# BCKDK regulates the TCA cycle through PDC to ensure embryonic development in the absence of PDK family

**DOI:** 10.1101/2020.03.22.002360

**Authors:** Lia Heinemann-Yerushalmi, Lital Bentovim, Neta Felsenthal, Ron Carmel Vinestock, Nofar Michaeli, Sharon Krief, Alon Silberman, Marina Cohen, Shifra Ben-Dor, Ori Brenner, Rebecca Haffner-Krausz, Maxim Itkin, Sergey Malitsky, Ayelet Erez, Elazar Zelzer

## Abstract

Pyruvate dehydrogenase kinases (PDK1-4) inhibit the TCA cycle by phosphorylating pyruvate dehydrogenase complex (PDC). Here, we show that the PDK family is dispensable for the survival of murine embryonic development and that BCKDK serves as a compensatory mechanism by inactivating PDC.

First, we knocked out all four *Pdk* genes one by one. Surprisingly, *Pdk* total KO embryos developed and were born in expected ratios, but died by postnatal day 4 due to hypoglycemia or ketoacidosis.

Finding that PDC was phosphorylated in these embryos suggested that another kinase compensates for the PDK family. Bioinformatic analysis implicated brunch chain ketoacid dehydrogenase kinase (*Bckdk*), a key regulator of branched chain amino acids (BCAA) catabolism. Indeed, knockout of *Bckdk* and the *Pdk* family led to loss of PDC phosphorylation, increment in PDC activity, elevation of Pyruvate flux into the TCA and early embryonic lethality. These findings reveal a new regulatory crosstalk hardwiring BCAA and glucose catabolic pathways, which feed the TCA cycle.

## Introduction

The pyruvate dehydrogenase complex (PDC) plays a major role as the gatekeeper that links glycolysis to the TCA cycle, maintaining metabolic balance and energy production through the rate-limiting and physiologically irreversible oxidative decarboxylation of pyruvate ^1^. Due to its importance, PDC regulation has been extensively studied since its discovery in the late 1960’s ^2^. The regulatory mechanism was shown to involve reversible phosphorylation by intrinsic regulatory enzymes, a family of pyruvate dehydrogenase kinases (PDK) 1-4 ^3^. These isoenzymes phosphorylate and inactivate PDC on three serine residues of its catalytic E1α subunit (PDH1a), namely S293 (site 1), S300 (site 2), and S232 (site 3). Studies of site specificity showed that all four PDKs phosphorylate site 1 and site 2, whereas site 3 is phosphorylated only by PDK1. Yet, phosphorylation of any of the three serine residues leads to complete inactivation of PDC activity ^4^.

PDK activity blocks the flux of pyruvate into the TCA cycle, which results in a metabolic shift to glycolysis for energy production. PDC activity is regulated by *Pdk* family in a short and long-term manner^5^. In the short term, several allosteric regulators can activate PDK according to the levels of end products, namely increased acetyl-CoA/coenzyme A ratio and reduced and oxidized nicotinamide adenine dinucleotide (NADH/NAD^+^) ratio^6^. In the long term, the amount of PDK protein is determined by different physiological conditions and in a tissue-specific manner ^7–10^. This involves transcriptional and translational regulation via hormonal regulators, such as estrogen receptor and glucocorticoid receptor^11^, and transcription factors such as oxygen sensor hypoxia inducible factor 1 subunit alpha (HIF1α) ^12–18^. HIF1α regulation of *Pdk* was shown to occur in cancer cells, which prefer to maintain anaerobic metabolism even in the presence of oxygen, a phenomenon known as the Warburg effect or aerobic glycolysis^19^. Here, the increase in glycolysis supplies intermediates for branching pathways that synthesize the macromolecules necessary for cell proliferation^20^. Based on these studies, *Pdk* family and specifically PDK1 became a specific target for anti-cancer drug development ^5,21^. In addition to its role in cancer, the HIF1-PDK axis is also vital for mammalian embryonic development^22–24^. For example, during endochondral bone formation, HIF1α is required for various processes in the hypoxic growth plate ^25–29^, one of which is to directly regulate the expression of *Pdk1* ^30^. Extensive studies on the involvement of the *Pdk* family in metabolic regulation have provided some indications for functional redundancy among its members. Cell culture experiments showed redundancy between PDK1 and PDK2 ^31^, whereas loss-of-function studies in mice lacking *Pdk2, Pdk4* or even both revealed no major effect ^31–33^. Nevertheless, the question of the necessity of the four PDK isoenzymes and the functional redundancy among them has yet to be addressed by a direct genetic approach.

In this study, we investigated the requirement of Pdk’s for embryonic development using the hypoxic growth plate as a model. Strikingly, mouse strains of double, triple and eventually quadruple KO, i.e. mice lacking all four *Pdk* genes, displayed normal skeletal development and were born in the expected Mendelian ratios. These results suggest that the entire *Pdk* family is dispensable for embryogenesis. Moreover, we show for the first time that in the absence of all PDK isoenzymes, PDC is still phosphorylated, implying the existence of a backup mechanism. Bioinformatic analysis implicated BCKDK, an enzyme that regulates the catabolism of branch chain amino acids, in this mechanism. The loss of PDC phosphorylation and consequent increases in PDC activity and pyruvate flux into the TCA cycles, which we observed in mice and cell lines lacking all *Pdk*’s and *Bckdk*, strongly support this possibility. Overall, we identify BCKDK regulation of PDC as a novel mechanism that can backup PDK family function to maintain embryonic development.

## RESULTS

### PDK1, PDK2 and PDK4 are dispensable for development and growth

To study in vivo the role of PDK family, we first focused on PDK1. Since the involvement of this isoenzyme in metabolic homeostasis was mostly studied in the context of cells under hypoxic conditions^14,15,34,35^, we chose the hypoxic growth plate during mouse bone development as a model^25,26,29,30,36^. Using the knockout-first allele method, we generated three mouse strains: floxed-*Pdk1, Pdk1-lacZ* knockout (KO) and *Pdk1* KO (Fig. S1A). Examination of the *Pdk1-lacZ* KO reporter line verified strong *Pdk1* expression in developing bones of E14.5 embryos, including long bones, vertebrae and facial bones (Fig. S1B-C). Surprisingly, both homozygous *Pdk1-lacZ* KO and *Pdk1* KO mice had no apparent bone phenotype during embryonic development, as confirmed by skeletal preparation (Fig. 1A,B) and histological sections (Fig. 1C,D). These mice produced viable and fertile colonies matching expected Mendelian ratios (Fig. 1G).

**Figure 1.**
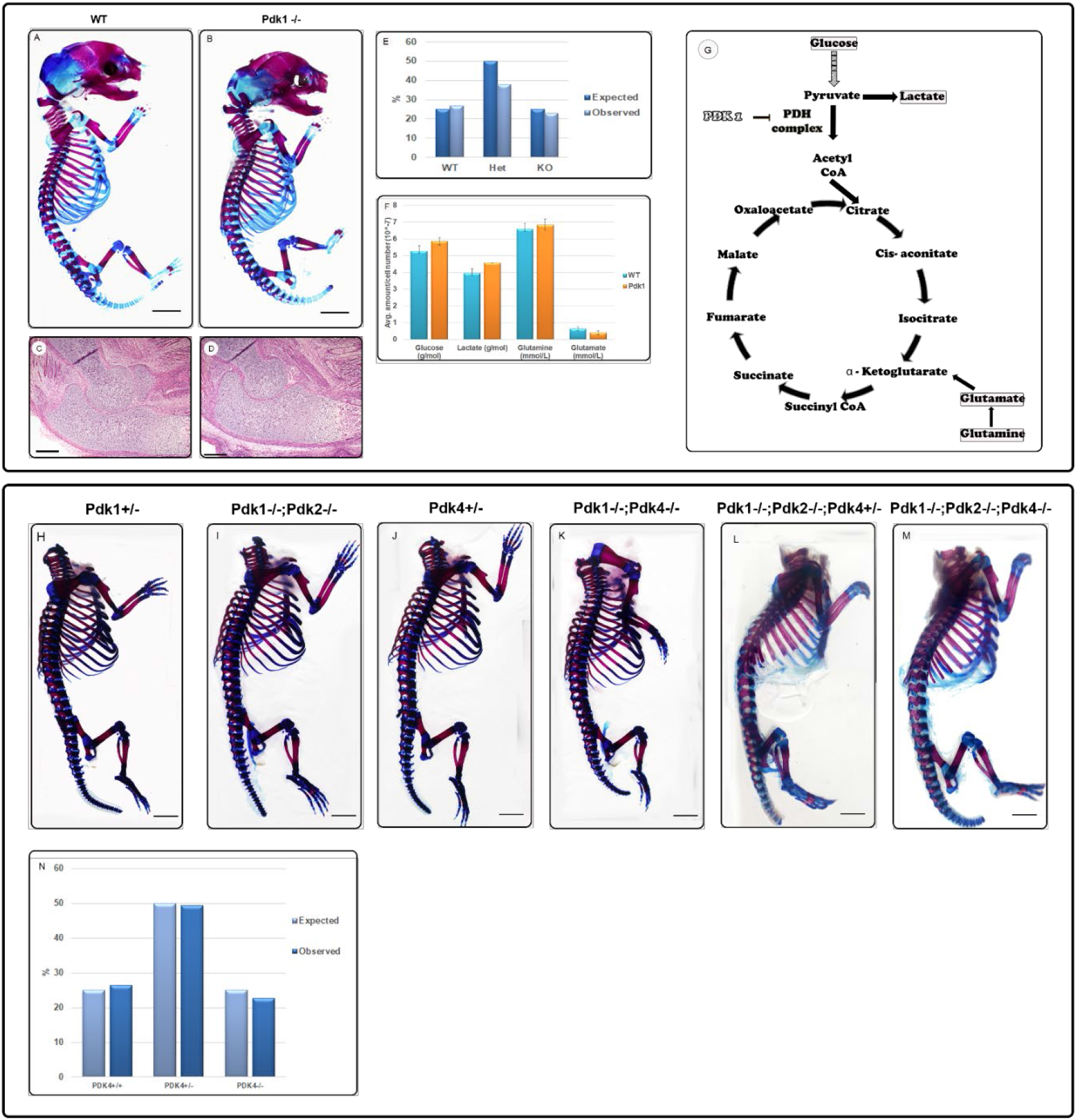
Deletion of single, double or triple *Pdk* genes in vivo does not affect embryonic development. (**A,B**) Skeletal preparations of control **(A)** and *Pdk1* KO **(B)** E17.5 embryos showing no skeletal phenotype (scale, 50 μm). (**C,D**) H&E staining of control and *Pdk1* KO E15.5 embryonic sections of olecranon and distal humerus growth plates (scale, 50 μm). **(E)** Graph showing expected Mendelian ratios of *Pdk1* KO progeny. **(F)** Graph showing similar absolute levels of glycolysis-TCA pathway-related metabolites in chondrocytes from E17.5 *Pdk1* KO and WT embryos (Nova analyzer, n=3). **(G)** Schematic illustration of the TCA cycle highlighting the relevant metabolites entry point to the cycle. **(H-M)** Skeletal preparations of control (*Pdk1^+/-^*, H), *Pdk1-Pdk2* dKO (I), control (*Pdk4^+/-^*, J) and *Pdk1-Pdk4* Dko (K), control (*Pdk1^-/-^Pdk2^-/-^ Pdk4^+/-^*,l) and *Pdk1-Pdk2-Pdk4* tKO (M) newborn pups. (H and I, J and K, and L and M are littermates; scale, 50 μm). **(N)** Graph showing expected Mendelian ratios in colonies of *Pdk* tKO mice.

To understand the lack of phenotype, we examined the levels of different metabolites related to the TCA and glycolysis pathways in growth plate chondrocytes from *Pdk1* KO and control mice. Chondrocytes from tibiofemoral growth plates were extracted and grown in cell culture, and the TCA and glycolysis intermediates were examined. Results showed similar levels of metabolites (glucose, glutamine and glutamate) in control and *Pdk1* KO chondrocytes, indicating that metabolic homeostasis in those cells is maintained. Moreover, we detected similar levels of lactate, suggesting that a metabolic shift to enhanced glycolysis occurred in the absence of *Pdk1* (Fig. 1F).

A plausible explanation for the lack of phenotype in the absence of *Pdk1* is that other PDK isoenzymes compensated for its activity. Null mutants for *Pdk2* and *Pdk4* were reported to develop normally and have viable and fertile colonies ^31,32^. Therefore, to examine the functional redundancy hypothesis, we generated double KO (dKO) and triple KO (tKO) mouse models lacking *Pdk1* and *Pdk2*, or *Pdk1* and *Pdk4*, or all three of these genes. The loss of these genes was verified by quantitative RT-PCR (qRT PCR) (Fig. S2A). Surprisingly, none of the combinations exhibited any bone phenotype (Fig. 1H-M). Moreover, *Pdk* tKO mice produced viable and fertile colonies matching expected Mendelian ratios (Fig. 1N). These outcomes indicated that PDK1, PDK2 and PDK4 are dispensable for bone and embryonic development.

### Loss of the *Pdk* family results in postnatal lethality due to hypoglycemia and ketoacidosis

The lack of phenotype in the *Pdk* tKO mice led us to study the only remaining candidate from the *Pdk* family, namely *Pdk3*. Recently, *Pdk3* was also shown to be a HIF1α target under hypoxic conditions ^18^. Thus, we hypothesized that *Pdk3* might be sufficient to maintain embryonic development. To test this hypothesis, we established a genetic model in which the expression of all four *Pdk* genes is deleted. Considering the possible embryonic lethality, we sought to delete *Pdk3* on the background of an intact *Pdk1* using *Pdk1^flox/flox^* allele combined with *Pdk2* and *Pdk4* KO. By crossing these mice with a specific Cre line, we could remove *Pdk1* in a tissue-specific manner and, thereby, prevent early lethality. For that purpose, we utilized the CRISPR/Cas9 method to target *Pdk3* gene on the genetic background of *Prx1-Cre;Pdk1^flox/null^ Pdk2^-/-^Pdk4^-/-^* mice and generated two independent lines. The first was the conditional quadruple knockout (cKO) *Prx1-Cre;Pdk1^flox/flox^Pdk2^-/-^Pdk3^-/-^Pdk4^-/-^* mice, which lack all four *Pdk* genes in *Prx1*-expressing limb mesenchyme lineages only. The second line was back-crossed to generate *Pdk1^-/-^ Pdk2^-/-^Pdk3^-/-^Pdk4^-/-^* quadruple KO mice (total KO) (Fig. 2A).

**Figure 2.**
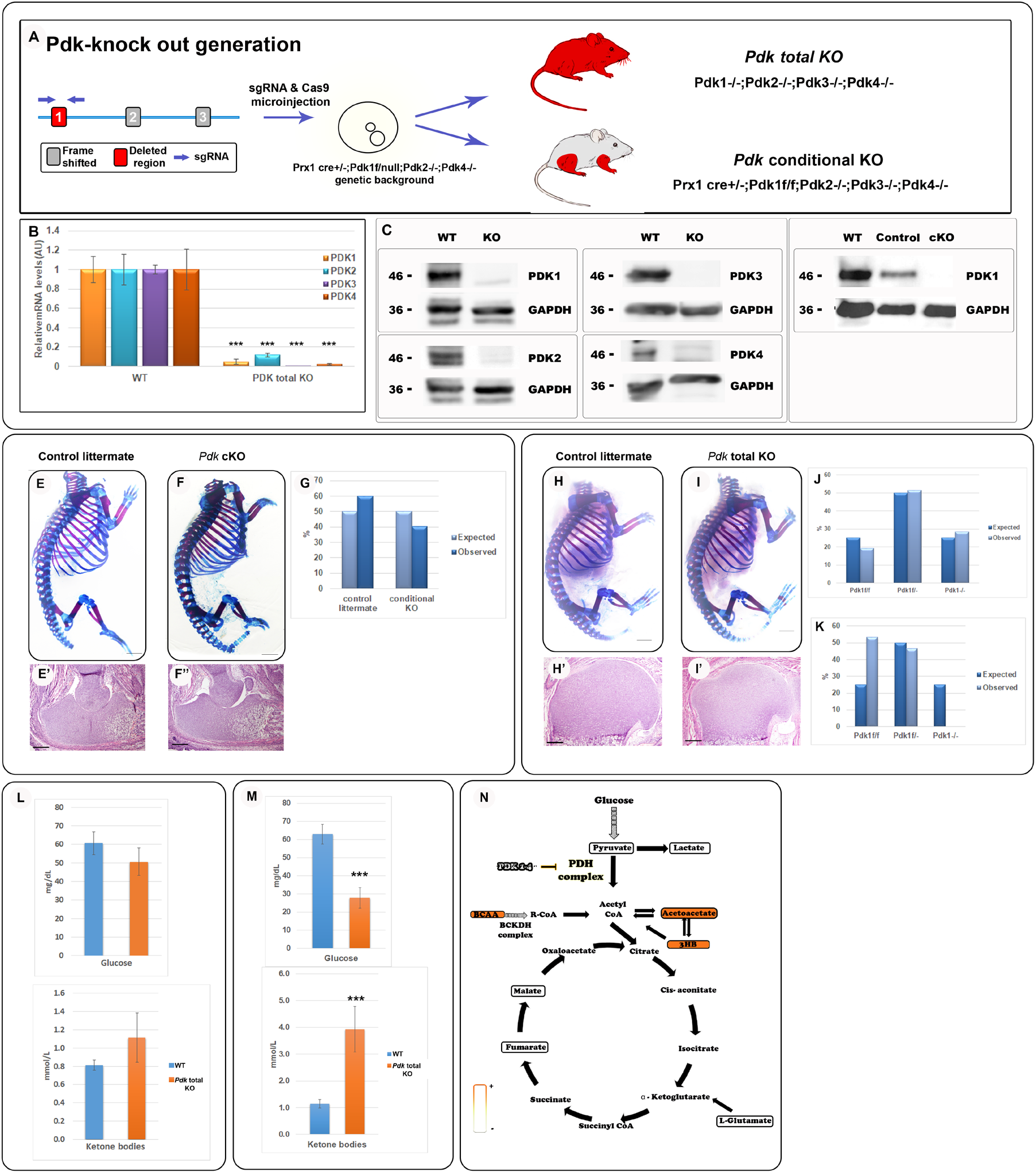
Deletion of the entire *Pdk* family results in postnatal lethality. (A) Schematic representation of *Pdk3* gene and the targeted site of CRISPR/Cas9-mediated deletion. This was followed by zygote injection to generate *Pdk* total KO mice or *Pdk* conditional KO in limb mesenchyme only, depending on the genetic background. (B) qRT-PCR of *Pdk1-4* mRNA confirms deletion of all four genes, as compare to WT (***, P<0.0001; n=4 for each genotype; data are normalized to *Tbp* and presented as mean±SD). (C) Protein expression analysis (western blot) of four PDK isoenzymes in heart samples derived from *Pdk* total KO or WT P0 pups shows complete deletion of the entire PDK family (representative images, n=3). (D) Western blotting of chondrocytes from *Pdk* cKO, control littermate and WT embryos shows deletion of PDK1 in the mutant (representative images, 3 biological repeats, n=5 for each sample). (E-F’) Skeletal preparations (E,F) and H&E-stained olecranon and distal humerus sections (E’,F’) from E18.5 *Prx1-Cre;Pdk1^f/f^;Pdk2^-/-^;Pdk3^-/-^;Pdk4^-/-^* embryos and control littermates (without Cre) show that the *Pdk* cKO embryos develop normally (scale, 100 μm). (G) Graph showing Mendelian ratios of genotypes in *Pdk* cKO progeny. (H-I’) Skeletal preparations at E18.5 (H-I) and H&E-stained sections (H’,I’) from P0 *Prx1-Cre;Pdk1^f/-^;Pdk2^-/-^;Pdk3^-/-^;Pdk4^-/-^* mice and control littermates (scale, 100 μm). (J) Graph showing Mendelian ratios of genotypes in E18.5 *Pdk* total KO embryos. (K) Graph showing non-Mendelian ratios of genotypes in *Pdk* total KO mature mice, as a result of loss of the total KO genotype. (L) Graphs showing in utero blood levels of glucose and 3HB, a ketone body, in *Pdk* total KO E18.5 embryos and P1 pups (M), compared to WT. Data are presented as mean ± SE (embryos: nWT=9, nKO=6; P1: nWT=9; nKO=4, from 3 independent litters; P<0.05, Student’s *t*-test). (N) Scheme of serum metabolic profiles of P1 *Pdk* total KO pups, as determined by LC-MS polar metabolite analysis. Higher levels are shown in orange and lower levels in yellow, as compared to WT levels (P<0.05; fold change >2, nWT=4; nKO=3).

Using qRT-PCR and western blot analysis on *Pdk* total KO mice, we verified deletion of all four *Pdk* genes at both mRNA and protein levels (Fig. 2B,C). Moreover, western blot analysis of chondrocytes derived from *Pdk* cKO growth plates showed deletion of PDK1 protein compared to cells from WT and control littermates, where *Prx1-Cre* was not present (Fig. 2D). The results indicated effective deletion of all four *Pdk* genes in both total KO and cKO mice. Surprisingly, skeletal preparation and histological sections from *Pdk* cKO mice revealed no bone phenotype (Fig. 2E-F’). Monitoring the colonies, we found that *Pdk* cKO mice produced viable and fertile colonies matching expected Mendelian ratios (Fig. 2G). Similarly, examination of bone development in skeletal preparation and histological sections of *Pdk* total KO embryos revealed no major skeletal phenotype (Fig. 2H-I’). However, although embryonic Mendelian ratios were observed as expected (Fig. 2J), this mouse strain did not produce viable offspring, as shown by the non-Mendelian ratios of mature mice (Fig. 2K).

Examination of newborn *Pdk* total KO pups revealed that they died between P0 and P4. To determine the cause of death, we analyzed histological sections from adrenal gland, brain, brown fat tissue, heart, kidney, liver, lung, small intestines and spinal cord of E18.5 embryos. The examination revealed no major anatomical abnormalities (Fig. S3).

Previous study showed that *Pdk* loss results in high utilization of glucose, followed by elevation of ketone bodies^33^. We therefore examined blood levels of glucose and ketone bodies as a possible cause of lethality. As observed in Fig. 2L-M, whereas at E18.5 no differences where observed between WT and *Pdk* total KO embryos, P1 *Pdk* total KO pups displayed significantly higher levels of 3-β-hydroxybutyrate (3HB), a common ketone body, and lower glucose levels as compared to WT pups (Fig. 2M).

Deeper metabolic profiling of serum composition in these pups showed significantly low levels of pyruvate and lactate together with a higher utilization of acetyl CoA, reflected by higher levels of the ketone bodies acetoacetate and 3HB, reduced production of more distal TCA metabolites, and high levels of branched chain amino acids (BCAA) (Fig. 2N). These results are consistent with the expected effect of *Pdk* loss ^33,37^, and suggest that the cause of the early postnatal death was ketoacidosis and hypoglycemia. Altogether, these results suggest that *Pdk* family is dispensable for embryonic development but is necessary postnatally to maintain energy metabolic balance.

### PDC phosphorylation is maintained upon PDK family loss of function

To gain molecular insight into the viability of *Pdk* total KO embryos, we studied the three phosphorylation sites on PDH1a subunit, which regulate PDC activity. First, we examined by western blot analysis PDC phosphorylation sites in chondrocytes isolated from growth plates of *Pdk* cKO embryos, which lack all *Pdk* genes in limb mesenchyme lineages. Interestingly, we found that one of the three sites, namely S300, was phosphorylated (Fig. 3A). To rule out the possibility that the phosphorylation of site S300 resulted from PDK1 contamination by surrounding tissues, such as skeletal muscles, we produced mouse embryonic fibroblasts (MEFs) from the total *Pdk* KO mice. As seen in Figure 3B, site S300 was phosphorylated in these MEFs as well. To strengthen the observation that the phosphorylation is maintained upon the loss of the *Pdk* family, we examined PDC phosphorylation in other tissues such as heart, lungs and kidney of *Pdk* total KO embryos. As seen in Figure S4, in all tested tissues, PDC was phosphorylated on at least one site in a tissue-specific manner. Altogether, these results suggest the existence of a backup mechanism that phosphorylates and inactivates PDC in the absence of all four PDKs.

**Figure 3.**
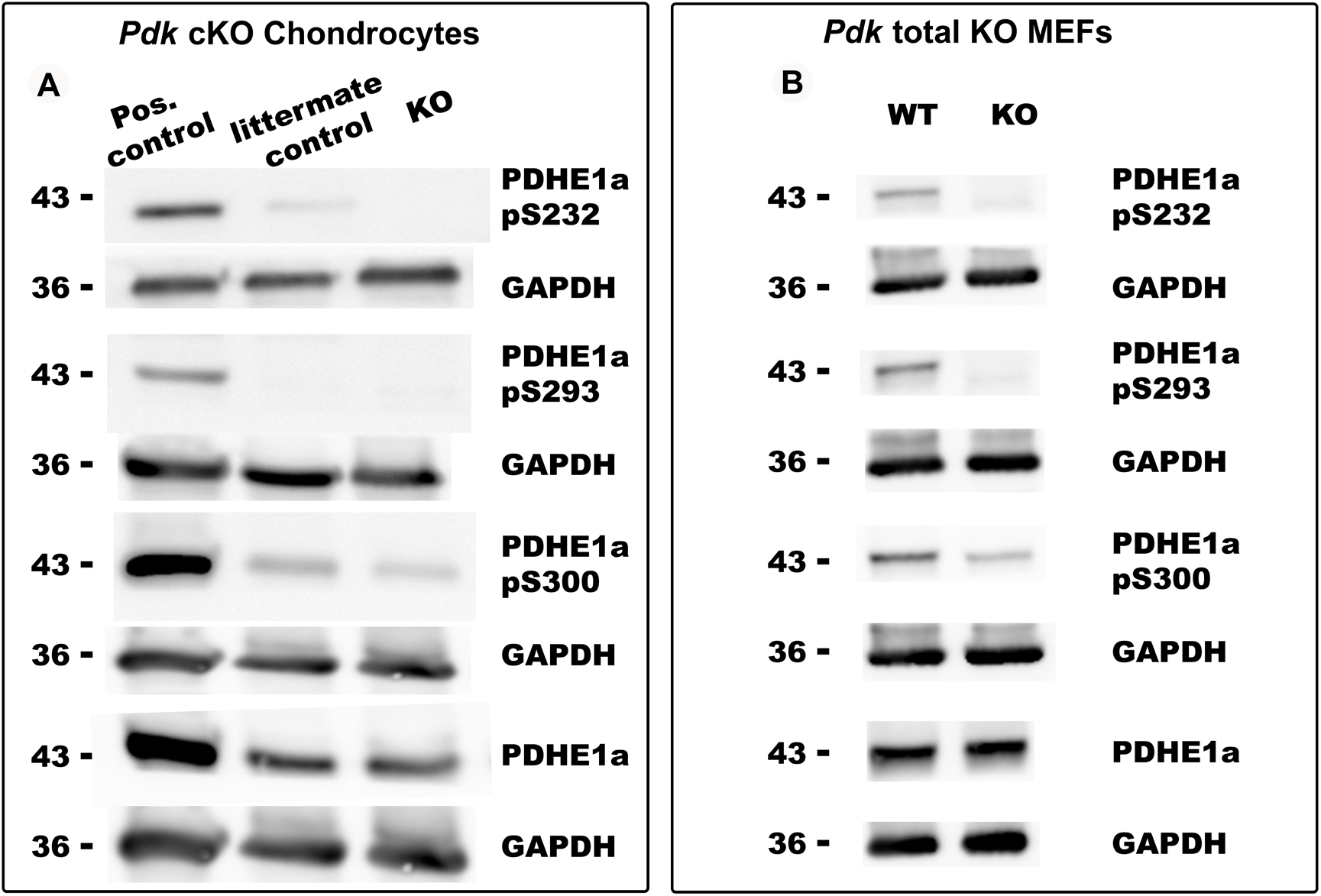
PDC phosphorylation is PDK-independent. Western blot analysis of the three PDC phosphorylation sites in chondrocytes extracted from growth plates of E17.5 *Pdk* cKO embryos (A; *Prx1-CrePdk1^flox/flox^Pdk2^-/-^Pdk3^-/-^Pdk4^-/-^*) and MEFs derived from *Pdk* total KO embryos (B), compared to WT. As controls, PDH1a and endogenous GAPDH total protein levels were measured. Each chondrocyte sample is a pool of 5 biological repeats; n=3 for all groups. Hearts of E17.5 WT embryos are shown as a positive control, littermate control were *Prx1-Cre*-negative. MEF samples, n=7.

### *Bckdk* is necessary for embryonic development in the absence of *Pdk* family

Our results suggested that another kinase, which is not a PDK, phosphorylates PDH1a subunit of PDC. To identify this kinase, we searched for suitable candidates using bioinformatic tools, such as Ensemble^38^ and GeneCards^39^. One candidate that emerged was branched chain ketoacid dehydrogenase kinase (*Bckdk*), which catalyzes the phosphorylation and inactivation of the branched-chain alpha-ketoacid dehydrogenase complex (BCKDC), the key regulatory enzyme of the valine, leucine and isoleucine catabolic pathways^40^. GenesLikeMe analysis^41^ predicted that *Bckdk* was a paralog of *Pdk1* and belonged to the same mitochondrial kinase proteins family, mainly by sequence and domain similarity (Fig. 4A). Moreover, protein-protein interaction analysis by STRING^42^ predicted BCKDK to interact with PDH1a subunit of PDC (Fig. 4B).

**Figure 4.**
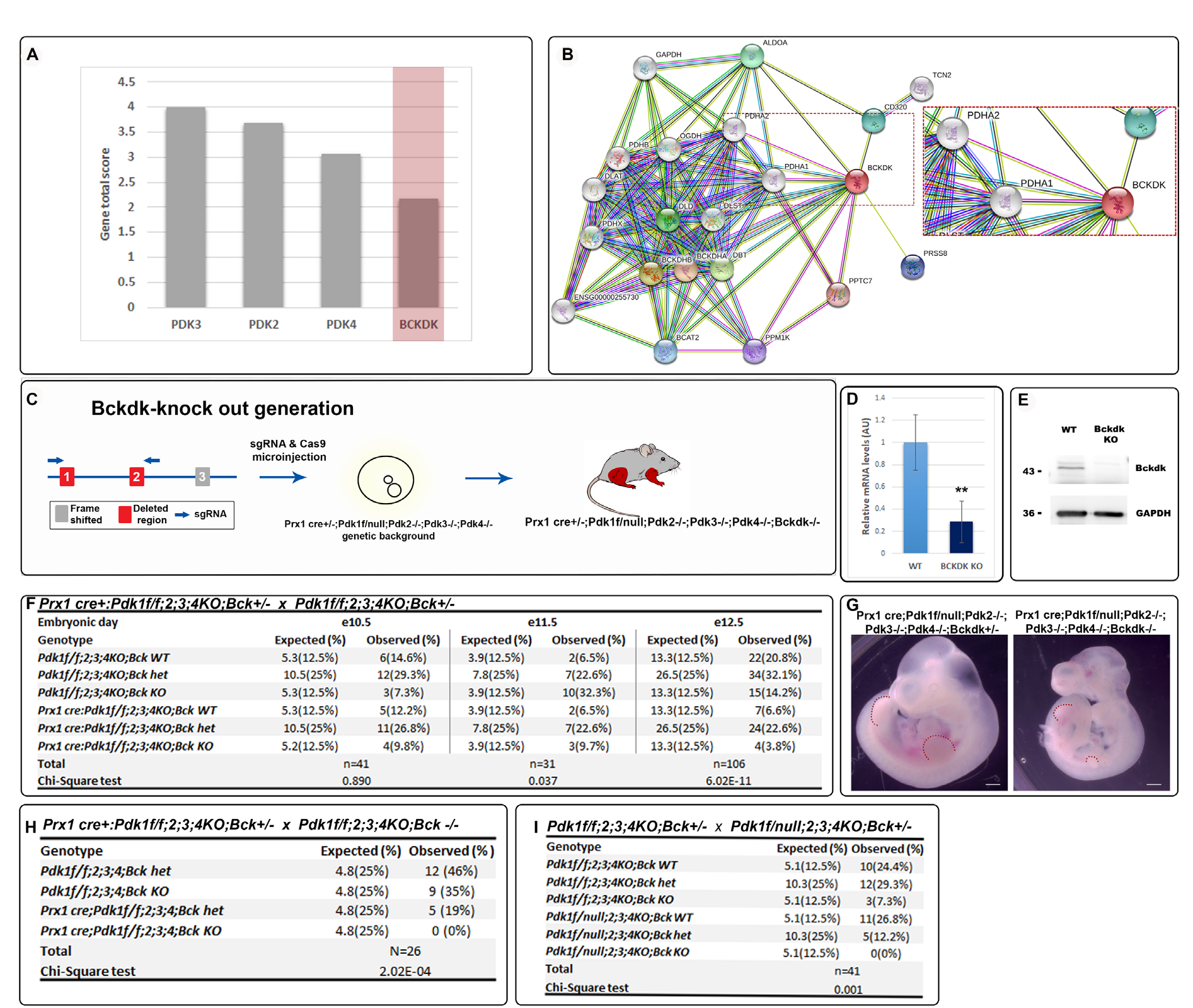
*Prx1-CrePdk cKO-Bckdk* KO mice display embryonic lethality. **(A)** GenesLikeMe analysis identifies *Bckdk* as a *Pdk1* paralog, along with the other *Pdk* family members (total score combines sequences, domains, super pathways, expression, compounds and gene ontology similarities). **(B)** STRING analysis predicts protein-protein interaction between BCKDK and PDH complex subunit PDH1a (string legend: yellow, textmining; black, co-expression; pink, experimentally determined). **(C)** Schematic representation of *Bckdk* gene and the targeted sites of CRISPR/Cas9-mediated deletion. This was followed by zygote injection to generate *Prx1-Cre-Pdk1^flox/null^ Pdk2k^-/-^Pdk3^-/-^4^-/-^Bckdk^-/-^* mouse, lacking *Pdk1* in limb mesenchyme lineages. **(D)** qRT-PCR of *Bckdk* mRNA in *Prx1-Cre-Pdk1^flox/null^ Pdk2^-/-^Pdk3^-/-^4^-/-^ Bckdk^-/-^* mice, as compare to WT (**, P<0.001; n=3 for each genotype; data normalized to *Tbp* and presented as mean±SD). **(E)** Protein expression analysis by western blotting shows complete deletion of BCKDK in *Pdk1^flox/flox^Pdk2^-/-^Pdk3^-/-^4^-/-^Bckdk^-/-^* MEF cells (n=3). **(F)** Expected Mendelian ratios and observed genotype distribution of *Prx1-Cre-Pdk1^floxflox^Pdk2^-/-^Pdk3^-/-^4^-/-^Bckdk^+/-^* crossed with *Pdk1^floxflox^Pdk2^-/-^ Pdk3^-/-^4^-/-^Bckdk^+/-^* embryos at E10.5 (n=41), E11.5 (n=31) and E12.5 (n=106). **(G)** Representative E10.5 *Prx1-Cre-Pdk1^floxflox^Pdk2^-/-^Pdk3^-/-^4^-/-^Bckdk^-/-^* embryo displays delayed development as compared to control *Bckdk* heterozygous littermate (dashed red lines indicate the limbs; scale, 50 μm). **(H)** Mendelian ratios and genotype distributions of *Prx1-Cre-Pdk1^floxflox^Pdk2^-/-^Pdk3^-/-^4^-/-^Bckdk^+/-^* crossed with *Pdk1^floxflox^Pdk2^-/-^Pdk3^-/-^4^-/-^Bckdk^-/-^* mice and **(I)** *Pdk1^flox/null^ Pdk2^-/-^Pdk3^-/-^4^-/-^Bckdk^-/-^* crossed with *Pdk1^flox/null^Pdk2^-/-^Pdk3^-/-^4^-/-^Bckdk^+/-^* mice (postnatal stage, after weaning). Data are presented as numbers and percentage of the total progeny, statistical significance was determined by Chi-square test.

Based on these observations, we hypothesized that BCKDK serves as the backup mechanism for PDK family during embryonic development by regulating PDC phosphorylation to inhibit its activity. To test this hypothesis, we sought to delete *Bckdk* on *Pdk* family KO genetic background. *Bckdk* KO mice were previously reported to be viable at neonatal stages and to display growth retardation only at three weeks of age^43^. To avoid embryonic lethality, we opted to delete *Bckdk* on the background of *Pdk1* cKO combined with *Pdk2-4* KO. Thus, we used the CRISPR/Cas9 method to generate a *Prx1-Cre-Pdk1^flox/null^Pdk2^-/-^Pdk3^-/-^4^-/-^Bckdk^-/-^* mouse strain (Fig. 4C). The deletion of *Bckdk* gene was verified at both mRNA and protein levels by qRT-PCR and western blotting, respectively (Fig. 4D,E). Crossing *Prx1-Cre-Pdk1^flox/null^Pdk2^-/-^Pdk3^-/-^4^-/-^Bckdk^+/-^* to *Pdk1^flox/null^Pdk2^-/-^Pdk3^-/-^4^-/-^Bckdk^+/-^* mice, we generated three strains: *Prx1-Cre-Pdk1^flox/flox^Pdk2^-/-^Pdk3^-/-^4^-/-^Bckdk^-/-^, Prx1-Cre-Pdk1^flox/null^Pdk2^-/-^Pdk3^-/-^4^-/-^Bckdk^-/-^*, and *Pdk1^flox/null^Pdk2^-/-^Pdk3^-/-^4^-/-^Bckdk^+/-^*.

First, we examined the viability of embryos lacking *Pdk1* in limb mesenchyme lineages and null for *Pdk2, Pdk3, Pdk4* and *Bckdk*. For that, we crossed *Prx1-Cre-Pdk1^flox/flox^ Pdk2^-/-^Pdk3^-/-^4^-/-^Bckdk^+/-^* mice with *Pdk1^flox/flox^ Pdk2^-/-^Pdk3^-/-^4^-/-^Bckdk^+/-^* and analyzed genotype distribution between E10.5 and E12.5, a stage at which *Prx1-Cre* is activated^44^ (Fig. 4F). Results showed that at E10.5 and E11.5, the numbers of *Prx1-Cre-Pdk1^flox/flox^ Pdk2^-/-^Pdk3^-/-^4^-/-^Bckdk^-/-^* embryos were close to the expected Mendelian ratios. Yet, some of these embryos displayed developmental retardation in comparison to control littermates (Fig. 4G). By contrast, at E12.5 the prevalence of this genetic combination was lower than expected, indicating embryonic lethality (Fig. 4F). Moreover, we failed to observe this combination postnatally (Fig. 4H).

To study genetic interaction between *Pdk1* and *Bckdk*, we crossed *Pdk1^flox/null^Pdk2^-/-^Pdk3^-/-^4^-/-^Bckdk^+/-^* mice with *Pdk1^flox/null^Pdk2^-/-^Pdk3^-/-^4^-/-^Bckdk^+/-^* mice. As seen in Figure 4I, in the presence of an intact *Pdk1* gene, all three combinations of *Bckdk*, namely WT, heterozygous and null, were observed. In contrast, heterozygosity of both *Pdk1* and *Bckdk* resulted in 50% reduction from the expected ratio, and no *Pdk1^flox/null^Pdk2^-/-^Pdk3^-/-^4^-/-^Bckdk^-/-^* offspring were observed, suggesting genetic interaction between these genes.

Together, these results clearly suggest that loss of *Bckdk* on *Pdk* family KO background results in embryonic lethality and strongly support the hypothesis that BCKDK compensates for the absence of PDK isoenzymes during development.

### BCKDK regulates PDC phosphorylation and activity as well as pyruvate flux into the TCA cycle

To further validate our hypothesis, we proceeded to test directly if *Bckdk* is necessary for the phosphorylation of PDC subunit PDH1a in the absence of all *Pdk* family members. To overcome the early embryonic lethality of mice lacking all *Pdk* genes and *Bckdk*, we used *Pdk1^flox/flox^Pdk2^-/-^Pdk3^-/-^Pdk4^-/-^Bckdk^-/-^* embryos to generate primary MEF cultures. Next, we infected these cells with adeno-Cre virus to ablate the expression of *Pdk1*, or with adeno-GFP as a control. As seen in Figure 5A, upon infection with adeno-Cre, concomitantly with the reduction in PDK1 expression, PDH1a phosphorylation on site S300 was reduced as compared to control cells; however, some phosphorylation was still noticeable. Since we observed low levels of PDK1 expression in adeno-Cre infected cells, we inhibit the activity of the remaining PDK1 by supplementing the culture with dichloroacetate (DCA), a PDK inhibitor^45^. As seen in Figure 5A, DCA nearly eliminated the phosphorylation of site S300. These results provide strong molecular evidence to the ability of BCKDK to regulate PDC by phosphorylation.

**Figure 5.**
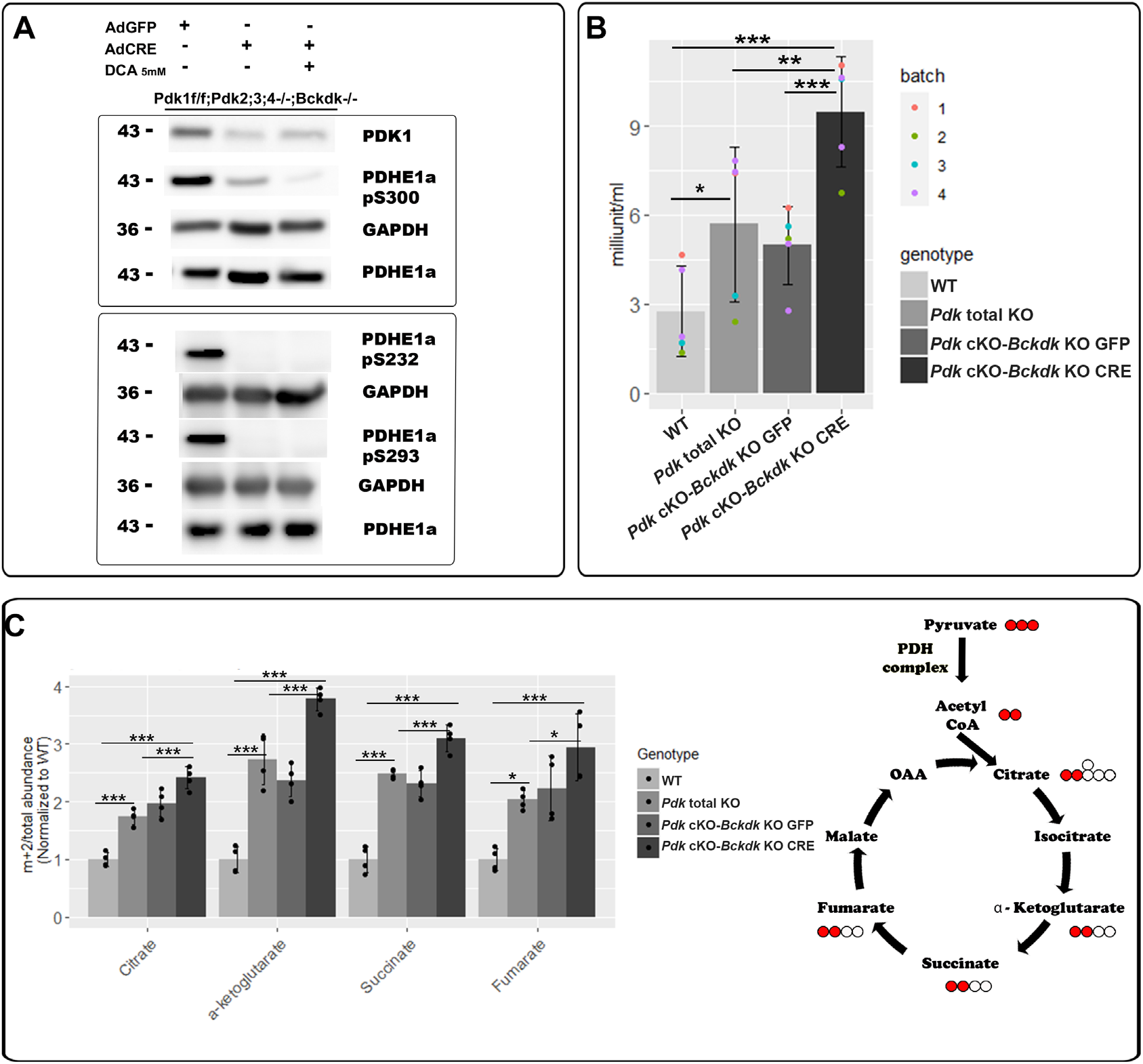
PDC phosphorylation and activity is BCKDK-dependent. **(A)** Western blot of PDK1 and of PDH1a phosphorylated sites pS300, the main active PDC site in MEFs, pS232 and pS293 in MEFs derived from *Pdk1^flox/flox^Pdk2^-/-^Pdk3^-/-^Pdk4^-/-^Bckdk^-/-^* mice infected with either Adeno-GFP as a control, or Adeno-CRE with or without addition of 5 mM DCA, to ablate PDK1 (representative images, n=6 from two different sets of experiments). **(B)** PDC activity assay in MEFs from *Pdk* total KO or *Pdk1^flox/flox^ Pdk2^-/-^Pdk3^-/-^Pdk4^-/-^Bckdk^-/-^* mice infected with either adeno-GFP (*Pdk* cKO-*Bckdk* KO GFP) or adeno-Cre (*Pdk cKO-Bckdk* KO CRE), and WT as a control (mean ± SD, *** P < 0.001, ** P < 0.01, * P < 0.05, n=5 for each genotype from 4 different experiments, ANOVA multiple comparison test). **(C)** Analysis of [3-^13^C]pyruvate flux into the TCA cycle using LC-MS. MEFs from *Pdk* total KO, *Pdk1^flox/flox^Pdk2^-/-^Pdk3^-/-^Pdk4^-/-^Bckdk^-/-^* or WT mice were infected with either adeno-GFP (*Pdk cKO-Bckdk* KO GFP) or adeno-Cre (*Pdk cKO-Bckdk* KO CRE). Graph shows the fraction of m+2 mass isotopomers of TCA cycle intermediates from the total abundance of these metabolites, normalized to WT levels. On the right is a schematics of the TCA cycle showing the labeled metabolites (mean ± SD, *** P < 0.001, ** P < 0.01, * P < 0.05, n=4 for each genotype, ANOVA multiple comparison test).

To demonstrate the biochemical effect of this regulation, we next quantified PDC activity in cells lacking *Pdk* family and in cells lacking both *Pdk* family and *Bckdk*. First, we examined PDC activity in MEFs from WT or *Pdk* total KO embryos. As seen in Fig. 5B, *Pdk* family loss resulted, as expected, in a significant two-fold increase in PDC activity, as compared to WT cells. Next, we examined PDC activity in our primary *Pdk1^flox/flox^Pdk2^-/-^Pdk3^-/-^Pdk4^-/-^Bckdk^-/-^* MEFs, which also lack *Bckdk*. Whereas in control MEFs infected with adeno-GFP virus (*Pdk* cKO-*Bckdk* KO GFP), which retain *Pdk1* expression, the increase in PDC activity was similar to the one observed in *Pdk* total KO cells, infection with adeno-Cre virus (*Pdk* cKO-*Bckdk* KO CRE) further increased the signal by 1.5 times relative to adeno-GFP infected MEFs and *Pdk* total KO-derived MEFs, and by 3 times relative to the WT.

Next, to provide direct metabolic evidence to the consequences of the observed changes in PDC activity, we examined pyruvate flux into the TCA cycle in these cell lines by performing ^13^C tracer experiments. Using labeled [3-^13^C]pyruvate, we measured the fraction of m+2 mass isotopomers of TCA cycle intermediates, namely citrate, α-ketoglutarate, succinate and fumarate, from the total abundance of these intermediates. As seen in Figure 5C, the fractions of labeled intermediates increased significantly in *Pdk* total KO MEFs relative to WT cells. Predictably, upon deletion of *Pdk* family and *Bckdk (Pdk cKO-Bckdk* KO CRE), the fractions of these metabolites further increased, as compared to both GFP control (*Pdk* cKO-*Bckdk* KO GFP) and *Pdk* total KO MEFs.

Collectively, these results provide strong molecular and biochemical evidence to the ability of BCKDK to regulate PDC phosphorylation and, thereby, activity.

## DISCUSSION

The ability of cells to adapt to different environmental conditions and maintain energetic homeostasis is one of the hallmarks of life. Using oxygen as the most common electron acceptor allows cells to maximize utilization of acetyl CoA driven from three catabolic pathways of glucose, amino acids and fatty acids feeding the TCA cycle. In this work, we discover a novel regulatory metabolic circuit between glucose and BCAA catabolic pathway. We found that BCKDK, a key regulator of BCAA catabolic pathway, regulates the activity of PDC, the gatekeeper of the glucose catabolic pathway. As the underlying mechanism, we show that BCKDK regulates PDC phosphorylation. This finding establish BCKDK as a backup mechanism for the PDK family in regulating PDC activity, which allows cellular metabolic balance to sustain embryonic development (Fig. 6).

**Figure 6.**
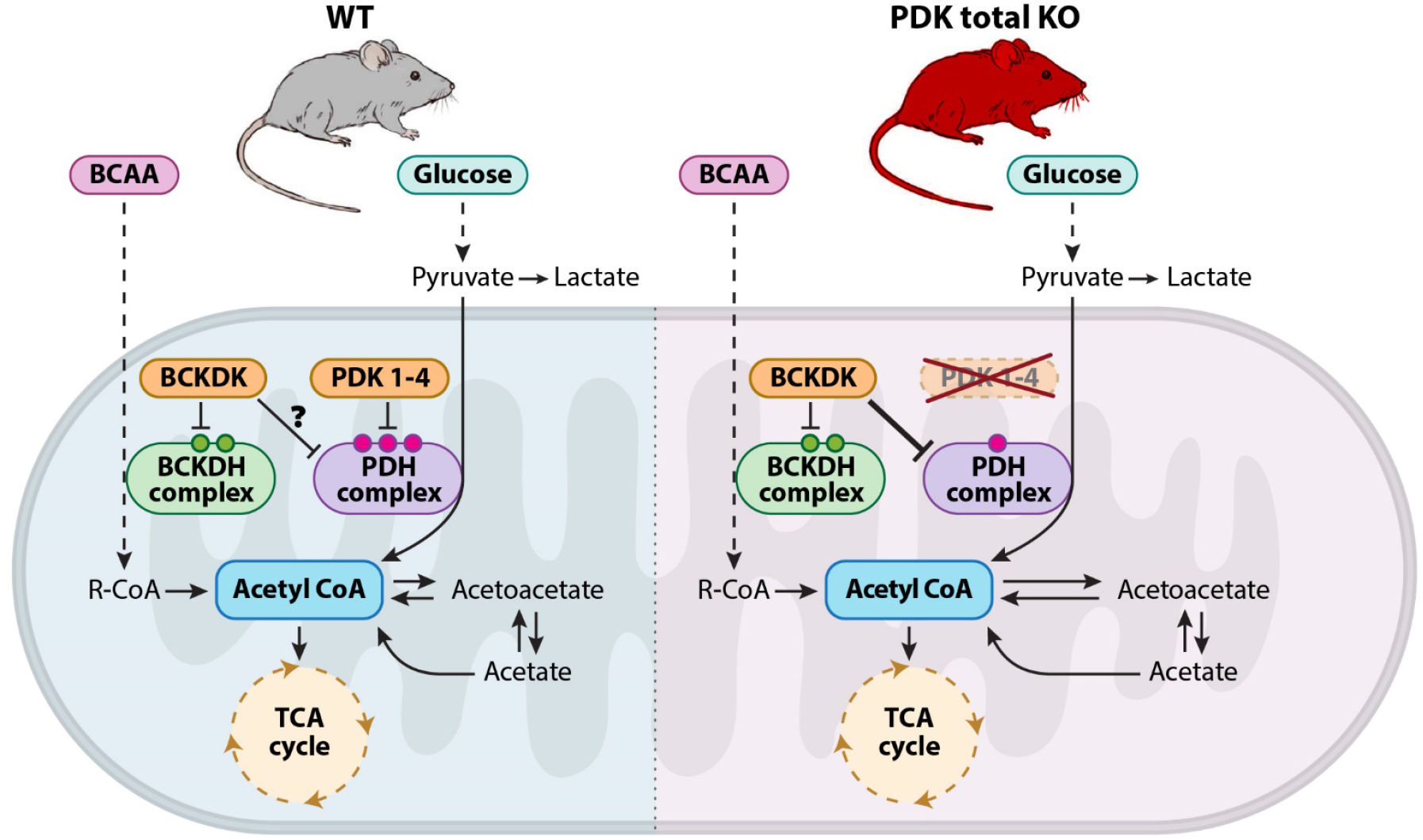
BCKDK compensates for *Pdk* loss of function via PDC phosphorylation. Cartoon depicting the interplay between glucose and BCAA catabolism in WT (left) versus *Pdk* total KO mice (right). In WT mice, PDC is inactivated by phosphorylation on three serine residues, whereas in the *Pdk* total KO mice, these phosphorylation sites are *Bckdk-* dependent. This novel level of regulation implicates BCKDK as a functional paralog of the PDK family, which may also act in the same way in the presence of PDKs.

BCAAs are a group of indispensable amino acids that includes leucine, isoleucine, and valine^43^. Their catabolism serves the cellular energetic balance by feeding the TCA cycle through a two-step degradation pathway of transamination to form branched chain α-keto acids (BCKAs), followed by oxidation and decarboxylation resulting in an irreversible synthesis of acetyl-CoA entering the TCA cycle^46^. The second step is crucial for degradation by BCKDC multienzyme. Therefore, BCKDC regulation is important to maintain proper levels of BCAAs and normal development^43^. Interestingly, PDC and BCKDC share many structural and enzymatic properties, as well as mechanism of regulation. Similarly to PDC, BCKDC is regulated by reversible phosphorylation of its E1 subunit on two serine sites (ser293 and ser303) mediated by its own native kinase, BCKDK. The activity of the entire BCKDC complex depends on BCKDK function, which is regulated by a feedback loop of substrate metabolites, e.g. BCAA blood levels ^47^.

Over the years, an interplay between the glucose and BCAA catabolic pathways has been established ^46^. Recently, it was shown that these pathways can regulate one another through feedback loops of nutrients blood-circulation, i.e, glucose and amino acids, to maintain energetic balance. Excessive or insufficient levels of glucose or BCAAs was shown to influence transcription of elements of the parallel pathway. For example, BCAA levels can improve glucose uptake by upregulation of glucose transporters ^48^, and a reciprocal effect was shown when high levels of glucose suppresses the expression of BCAA degradation enzymes.^49^. Our discovery that BCKDK can phosphorylate PDC uncovers a new level of regulation that hardwires glucose and BCAA degradation pathways through direct enzymatic regulation and not by a feedback loop of circulating nutrients.

From an evolutionary viewpoint, Oppenheim et. al ^50^ recently showed that in some parasites lacking mitochondrial PDC, acetyl-CoA is converted from pyruvate by BCKDC, which fulfills PDC function. Therein, it was suggested that BCKDC evolved earlier than PDC. Although the ancestral gene of the regulators of these complexes, i.e. *Pdk* and *Bckdk*, is yet to be discovered, sequence similarities between *Pdk* and *Bckdk* has placed them as paralogs belonging to the same mitochondrial kinase family^51–54^. Here, we provide strong in vivo evidence suggesting that *Bckdk* is not only a paralog, but also a functional paralog of the *Pdk* family. One interesting question that remains is whether BCKDK acts only as a backup mechanism for the loss of the PDK family, or is it a part of PDC regulation in normal conditions as well. Another intriguing question that is raised by the observed functional compensation is whether PDKs can compensate for the absence of BCKDK. This assumption makes sense considering that *Bckdk* KO mice are viable and exhibit growth retardation only at three weeks of age, due to lack of BCKDC regulation^43^.

Another interesting point emerging from our finding that BCKDK can regulate PDC is the tissue-specific expression of PDK isoenzymes^52^. Moreover, each isoenzyme was shown to be regulated by different physiological conditions^5,9,10^. By contrast, several studies support the possibility of functional redundancy among PDK family members, for example, between PDK1 and PDK2 in muscle tissue in vitro^31^ and between PDK2 and PDK4 in vivo^33^. Our failure to identify major developmental phenotype in embryos that lost different *Pdk* genes increases the scope of known functional redundancy among these four isoenzymes. The sequential targeting of these genes demonstrates that each PDK can compensate for the loss of the other family members. Another level of this complex regulation relates to the tightly controlled rate of PDC phosphorylation, which is maintained by the balance between the activity of PDK and pyruvate dehydrogenase phosphatases (PDPs). The latter are two enzymes that reverse PDK activity and thereby reactivate PDC^55^. Given the importance of this balance, it is interesting to see whether the activity levels of PDKs and PDPs are regulated such that a reduction in PDK activity would lead to a concomitant reduction in PDP activity. Such coordination might contribute to the maintenance of PDC phosphorylation in the absence of PDK family. Overall, PDC regulation by kinase versus phosphatase activity should be further studied under different physiological conditions.

On that note, hypoxic conditions have long been associated with pathologies, such as tissue ischemia, inflammation and cancer^56^. However, hypoxic microenvironments were shown to be essential for mammalian embryonic development^24^. In either state, HIF1 transcription factor initiates a key molecular response that maintains O2 homeostasis and metabolic balance^14,22^. Specifically, the HIF1α-PDK axis is considered a major regulator of cell adaptation to hypoxia by both controlling energy production and decreasing mitochondrial oxygen consumption^12–14,16,18,57^. However, our results of normal development in the absence of one side of this axis, namely all PDKs, question its significance. There are three possibilities to reconcile our findings with the common view. One is that HIF1α activation of a wide range of downstream target genes is adaptive enough to support metabolic energy balance under hypoxia even without inactivation of PDC. For example, by activating pyruvate kinase isoform M2 (PKM2), which diverts pyruvate to the pentose phosphate pathway ^18^. Second, it is possible that by activating another, yet undescribed mechanism, HIF1α compensates for the absence of PDK family. In that context, the possibility that HIF1α can regulate *Bckdk* should be considered. Finally, it is possible that the involvement of HIF1α-PDK axis is restricted to the development or function of specific organs, which are dispensable for embryonic development. In support of this explanation is our finding that mice lacking all *Pdk* genes in limb mesenchyme developed normally, whereas total *Pdk* KO mice died at early neonatal stages. The metabolic abnormalities in these newborn total KO mice strongly imply that the affected organs are involved in physiological regulation of the organism, which is critical for its survival postnatally but not during embryogenesis. It is also possible that during development, the consequences of metabolic abnormalities, such as ketone body accumulation and hypoglycemia, are buffered by the safe environment of the placenta^58^. Obviously, the different outcomes of embryonic development and postnatal growth of *Pdk* total KO mice does not exclude the possibility that these two processes have different metabolic requirements. In that case, it is possible that BCKDK could backup for PDKs only during the embryonic period.

To conclude, our findings provide new insight into the functional redundancy among PDK family members in modulating PDC activity to maintain energy production and cell survival, while revealing the existence of a previously unknown backup mechanism, which places *Bckdk* as a functional paralog of *Pdk*. This backup mechanism unveils a new level of regulatory crosstalk between two central metabolic pathways that feed the TCA cycle. The finding of this backup mechanism may promote the development of new therapeutic strategies for complex diseases involving changes in the metabolic state of cells, such as cancer, diabetes and many other metabolic diseases.

## Author contribution

Experiment Design, L.H-Y, L.B, and E.Z; Experiments, L.H-Y, L.B, N.F, R-C.V, N.M, S.K, A.S, M.C, S.B-D.O.B, R.H-K, M.I and S.M; Intellectual Contributions, E.Z, A.E, L.H-Y, L.B, N.F, R-C.V; Manuscript Writing, L.H-Y and E.Z with input from all authors.

## Acknowledgments

We thank Nitzan Konstantin for expert editorial assistance, Dr. Robert Harris from Indiana University, who kindly provided us with PDK2 and PDK4 mice, Dr. Nicola Brunetti-Pierri from the Telethon Institute of Genetics and Medicine for his support and advice, Dr. Ron Rotkoff for his help with statistical analysis, Dr. Tsviya Olender for bioinformatics advice, and Neria Sharabi from the Department of Veterinary Resources for his help with mouse maintenance. Special thanks to all members of the Zelzer Laboratory for encouragement and advice.

## STAR Methods

### RESOURCE AVAILABILITY

#### Lead Contact

Further information and requests for resources and reagents should be directed to and will be fulfilled by the Lead Contact, Elazar Zelzer

#### Materials Availability

All unique reagents generated in this study are available from the Lead Contact.

#### Data and Code Availability

The data reported in this study are available from the Lead Contact.

### EXPERIMENTAL MODEL AND SUBJECT DETAILS

#### Mouse lines

*Pdk2* KO (Vandenboom et al., 2011) and *Pdk4* KO mice (Ho Jeoung et al., 2006) were previously described. The *Pdk1* KO, floxed-*Pdk1* and *Pdk1-lacZ* mice were generated as follows: Embryonic stem cells with a gene trap insertion (KO-first allele) in the *Pdk1* gene were obtained from the European Conditional Mouse Mutagenesis program (EUCOMM) (Pettitt et al., 2013). *Pdk1* KO mice were crossed with PGK-Cre mice (Lallemand, Luria, Haffner-Krausz, & Lonai, 1998) to generate the null *Pdk1-lacZ* mice. Alternatively, *Pdk1* KO mice were crossed with *Rosa26-FLPe* mice (Jackson laboratory, 016226) to generate the floxed-*Pdk1* mice. Genotyping of *Pdk1* KO, *Pdk1-lacZ* and floxed-*Pdk1* mice was performed by PCR (primer sequences are shown in Supplementary Table S1).

Generation of double and triple KO combinations of *Pdk1, Pdk2* and *Pdk4* was done by strain breeding. *Pdk3* KO and *Bckdk* KO were generated as follows: Cas9 plasmid and plasmids encoding guide RNAs were designed and optimized for the best guides using several CRISPR designing tools, including the MIT CRISPR design tool (Hsu et al., 2013) and sgRNA Designer, Rule Sets 1 and 2 (Doench et al., 2016, 2014), in both the original sites and later in the Benchling implementations (www.benchling.com), SSC (Xu et al., 2015), and sgRNAscorer (Chari, Moosburner, & Church, 2015), in their websites. The following oligos were used for construction of gRNA: Pdk3: 5’-CGGGTTGGGGAGGTCTAGAG-3’ (upstream of the putative TSS) and 5’-ATTCCGTGAGAAGCTCCGGG-3’ (downstream of the ATG in the first exon), total deletion of 560 bp (location X:93831603-93832603). For Bckdk: 5’-CCCGCGCGATGTTACAGCCG-3’ (an upstream guide around the TSS) and 5’-CTCTACATGGTGTGTATCGG-3’ (downstream of the ATG in the second exon), total deletion of 884bp (location chr7:127903907-127905206). In vitro transcribed Cas9 RNA (100 ng ml 1) and sgRNA(50 ng ml 1) were injected into one-cell stage fertilized embryos isolated from superovulated *Prx1-Cre;Pdk1^flox/-^ Pdk2^-/-^Pdk4^-/-^* or *Prx1-Cre;Pdk1^flox/-^Pdk2^-/-^Pdk3^-/-^Pdk4^-/-^* mice mated with males of the same mutation to generate *Pdk* total KO or *Pdk* cKO-*Bckdk* KO mice, respectively. Injected embryos were transferred into the oviducts of pseudopregnant ICR females as previously described (Jaenisch et al., 2013). Genomic DNA from treated embryos was analyzed by PCR primers designed to verify the deleted sequence of each gene; primer sequences are shown in Supplementary Table S1.

Staff and veterinary personnel monitored all mouse strains daily for health and activity. Mice were given ad libitum access to water and standard mouse chow with 12-hr light/dark cycles. All animal procedures were approved by the Institutional Animal Care and Use Committee and performed in strict adherence to Weizmann Institute Animal Care and Use guidelines, following the NIH, European Commission, and Israeli guidelines. In all timed pregnancies, plug date was defined as E0.5. For harvesting of embryos, timed-pregnant female mice were sacrificed by CO2 intoxication. The gravid uterus was dissected out and suspended in a bath of ice-cold PBS and the embryos were harvested. Tail genomic DNA was used for genotyping. All procedures and treatments are described as in Method Details. None of the mice was involved in any previous procedures prior to the study.

#### Primary cultures

For chondrocyte primary culture, hindlimb tibiofemoral growth plates of E17.5 *Pdk1* KO, *Prx1-Cre;Pdk1^flox/-^Pdk2^-/-^Pdk3^-/-^Pdk4^-/-^* or WT embryos were dissected and soft tissue, skin and particularly muscles were removed. The growth plates were dissected and placed in DMEM 4500 mg/l glucose (Thermofisher) with 1% pen-strep solution (Biological Industries). Growth plates were digested in trypsin containing 0.25% EDTA (Biological Industries) for 30 minutes at 37°C and in 1 mg/ml collagenase type V (Sigma) in DMEM 4500 mg/l glucose with 1% pen-strep solution for 2 hours. Chondrocytes were plated at a density of 125×103 cells/ml and grown under normoxia in monolayer cultures in high glucose DMEM supplemented with 10% fetal bovine serum (FBS, Biological Industries) and 1% pen-strep solution for 5 days until confluency (>90%).

Mouse embryonic fibroblasts (MEFs) were extracted from E12.5-E14.5 *Pdk* total KO, *Prx1-Cre;Pdk1^flox/flox^Pdk2^-/-^Pdk3^-/-^Pdk4^-/-^* or ICR (WT) embryos, and cultured under normoxic conditions in DMEM (Gibco), 20% fetal calf serum, 1% L-glutamine,1% MEM-non-essential amino acid (Biological Industries), 1% penicillin/ streptavidin and sodium pyruvate. At passage 3 or 4, cells were harvested for either qRT-PCR, western blot analysis or viral infection. All procedures and treatments are described in Method Details.

### METHOD DETAILS

#### Viral infection

For adeno-viral infection, MEFs were infected with 350 viral particles/cell of Ad5CMVeGFP or Ad5CMVCre-eGFP virus (Gene Transfer Vector Core, University of Iowa). MEFs were plated at a density of 500×10^3^ cells in 6-cm plates with the same growth medium containing the AdCMV virus for 24 h, when medium was added. Medium was replaced after 48 h and cells were harvested for western blot analysis or metabolic LC-MS/MS analysis 5 days post-infection. During all experiments, medium was changed daily.

#### Histology

For hematoxylin and eosin staining (H&E), embryos were fixed overnight in 4% paraformaldehyde (PFA) -phosphate-buffered saline (PBS), decalcified in a solution containing equal parts of 0.5 M ethylenediaminetetraacetic acid (EDTA; pH 7.4) and 4% PFA in PBS overnight, dehydrated to 100% ethanol, embedded in paraffin and sectioned at a thickness of 7 μm. For pathological examination, sections of the entire embryo were made. H&E staining was performed following standard protocols.

#### X-gal staining

Whole-mount X-gal staining was performed as described previously (Eshkar-oren et al., 2009). In short, freshly dissected tissue was fixed in 4% PFA/PBS, rinsed in a solution containing 5 mM EGTA, 0.01% deoxycholate, 0.02% NP40 and 2 mM MgCl_2_, and then stained in a solution containing 5 mM K_3_Fe(CN)_6_, 5 mM EGTA, 0.01% deoxycholate, 0.02% NP40, 2 mM MgCl_2_ and 1 mg/ml X-gal. The tissue was cleared in 0.3% KOH for better visualization.

#### Skeletal preparation

Cartilage and bones in whole mouse embryos or newborn pups were visualized after staining with Alcian Blue and Alizarin Red S (Sigma) and clarification of soft tissue with potassium hydroxide (Mcleod, 1980).

#### Glucose, glutamine, glutamate and lactate measurements

Chondrocytes were plated as described above. Upon reaching 90% confluency, cells were washed and incubated in a glucose- and glutamine-free DMEM medium supplemented with 10% dialyzed serum, 4 mM L-glutamine 10 mM glucose, for 12 h. Subsequently, 500 μL medium from the cell culture was collected, briefly centrifuged and directly injected into the Nova chemical analyzer. Background metabolite measurements from cell-free culture were subtracted, and results were normalized to cell number.

#### RNA isolation and quantitative real-time (qRT-) PCR

Total RNA was purified from either MEFs or chondrocyte primary culture using the RNeasy Kit (Qiagen). Reverse transcription was performed with High Capacity Reverse Transcription Kit (Applied Biosystems) according to the manufacturer’s protocol. qRT-PCR was performed using Fast SYBR Green master mix (Applied Biosystems) on the StepOnePlus machine (Applied Biosystems). Values were calculated using the StepOne software version 2.2, according to the relative standard curve method. Ct values were normalized to TATA-box binding protein (Tbp) or 18S rRNA. Statistical significance was determined by Student’s *t*-test as P<0.05.

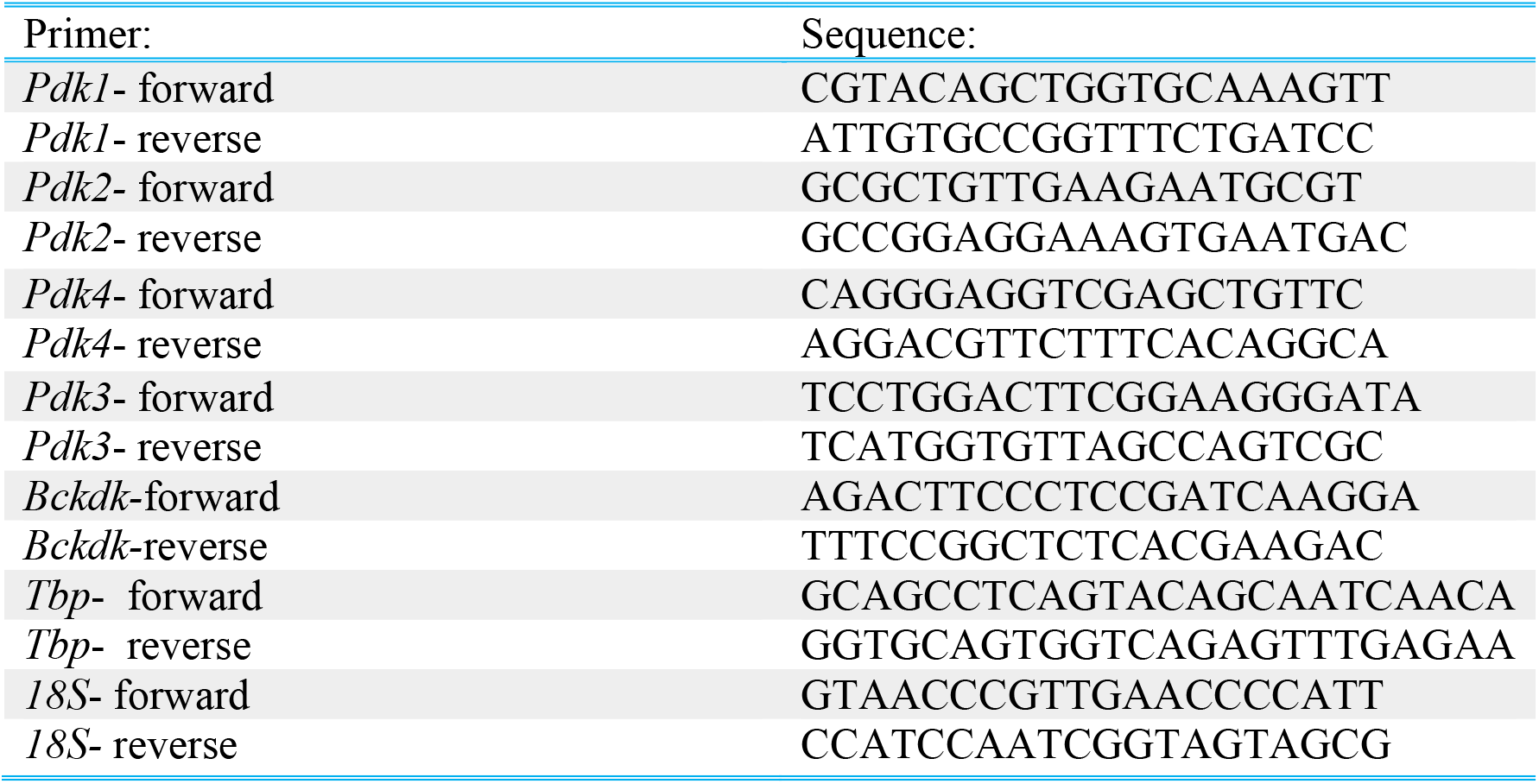

#### Western blot analysis

For western blotting, protein was extracted using RIPA (150mM NaCl, 1% NP40, 0.5% Deoxycholate, 0.1% sodium dodecyl sulfate (SDS), 50mM Tris pH 8) supplemented with PI (1:100) and phosphatase inhibitors (1:20), homogenized using Motor Cordless (Kimble) and centrifuged at 10,000 rpm, 4 °C for 10 min. Supernatant was collected and used for the following procedures. In the case of heart, kidney and lungs extracts were supplemented by sonication (SONICS, Vibra-Cell™) post-homogenization. Protein concentration was determined by Pierce protein BCA assay kit (Cyanagen). 40 μg protein extracts were denatured by boiling in × 5 sample buffer (60 mM Tris-HCl (pH 6.8), 25% glycerol, 2% SDS, 14.4 mM β-mercaptoethanol, 0.1% bromophenol blue) for 5 min, resolved by 10% SDS-PAGE, 100-120 V, for 70 min and transferred to nitrocellulose membrane (Whatmann, 10401383) at 250 mA for 70 min. Membranes were stained with Ponceau (Sigma-Aldrich, P7170) and blocked for 60 min at room temperature (RT) with 5% milk-0.05% TWEEN-20. Next, membrane was incubated rocking with primary antibodies in Antibody solution (5% BSA fraction V, 5% sodium azide, 0.05% PBST) overnight at 4 °C. Membranes were washed 3 × 5 min at RT with 0.05% PBST and incubated for 60 min with horseradish peroxidase (HRP)-conjugated secondary antibodies. Membranes were washed 3× 5 min, processed with EZ-ECL Chemiluminescence detection kit for HRP (Biological Industries, 20-500-120) and visualized by ImageQuant™ LAS 4000 (GE Healthcare Life Sciences). Densitometry values were normalized to GAPDH in the same lane.

#### PDH activity assay

PDH Activity Assay Kit (Sigma) was preformed according to the manufacturer’s instruction. In short, protein was extracted from ~10^6^ MEF cells, homogenized using Motor Cordless (Kimble) in 100 μL of ice-cold PDH Assay Buffer and centrifuged at 10,000 rpm, 4 °C for 5 min. Supernatant was collected and used immediately (20 ul per well). Colorimetric reads were done by Infinite® 200 PRO NanoQuant (Tecan), samples were normalized to cell count.

#### Ketone bodies and glucose blood levels measurements

3-beta-hydroxybutyrate (3HB), a prevalent ketone body, and glucose were measured by Freestyle Optium Neo machine (GeffenMedical) and Performa blood meter (ACCU-CHECK®), respectively, using a drop of blood from P1 *Pdk* total KO pups.

#### Bioinformatic analysis of potential Pdk paralogs

GenesLikeMe paralog analysis (https://www.genecards.org/) for *Pdk1* was performed as previously described (Stelzer G, Inger A, Olender T, Iny-Stein T, Dalah I, Harel A, Safran M, 2009). Protein-protein interaction analysis was done using STRING (Szklarczyk et al., 2017).

#### Genotype distribution analysis

All genotype ratios were analyzed by monitoring colony progeny (4-6 months) or embryos, using tail PCR genotyping followed by Chi-square test. n values are given in the figure legends.

#### Metabolite extraction

For serum metabolite composition analysis, total blood volume (about 50 μl) from P1 *Pdk* total KO pups were collected into serum separation tubes (MiniCollect Tubes, greiner bio-one) and centrifuged at 1000 rpm for 5 min at RT. Serum was collected and immediate frozen in safe-lock Eppendorf tubes in liquid nitrogen. For labeled metabolites, MEF primary culture were grown and infected as described. 5 days post-infection growth medium was removed, cells were washed with pre-warmed PBS and medium was replaced with DMEM free of L-Glutamine, Glucose and Sodium Pyruvate (Biological Industries) supplemented with 2.5 mM glucose, 2.5 mM [3-^13^C]pyruvate, 2 mM l-glutamine and MEM-eagle non-essential amino acids X1, and incubated at 37 °C, for 1.5 h. Then, medium was removed and cells were washed twice with 2 ml 0.9% ice-cold saline. Cells were scraped from plates with 500 μl of ice-cold methanol:DDW (1:1, v:v) containing C13 and N15 labeled amino acid mix (Sigma-Aldrich) as internal standards; a total volume of 1000 μl from 2X plates per sample (about 2X10^6 cells) were collected into safe-lock Eppendorf tubes. Samples underwent three freeze-thaw cycles in liquid nitrogen −37°C bath sonicated in ice-bath for 30 min, and vortexed each 10 min. Then, the samples were centrifuged for 15 min at maximum speed (14000 rpm) at 4°C. Supernatant (800 μL) was transferred to another Eppendorf tube, dried for 1 h in speedvac and lyophilized. The dry pellet was re-suspended in 100 μL methanol:DDW (1:1, v:v) and centrifuged twice for 15 min at maximum speed (14000 rpm) at 4°C. Then, the supernatant was transferred into vials for injection.

#### LC-MS polar metabolites analysis

Metabolic profiling of polar phase was done as described by Zheng et al. (2015) with minor modifications. Briefly, analysis was performed using Acquity I class UPLC System (Waters) combined with Exactive™ Plus Orbitrap Mass Spectrometer (Thermo Scientific™), which was operated in a negative ionization mode. The LC separation was done using the SeQuant Zic-pHilic (150 mm × 2.1 mm) with the SeQuant guard column (20 mm × 2.1 mm) (Merck). The Mobile phase A consisted of acetonitrile and Mobile phase B consisted of 20 mM ammonium carbonate plus 0.1% ammonia hydroxide in water. The flow rate was kept at 200 μl min-1 and gradient as follow: 0 −2 min 75% of B, 17 min 12.5% of B, 17.1 min 25% of B, 19 min 25% of B, 19.1min 75% of B, 19 min 75% of B.

#### Polar metabolites data analysis

The data were collected using Xcalibur 4.15 and processed using Qual Browser (Thermo Scientific™). Compounds were identified by retention time and fragments and were verified using in-house mass spectra library. Metabolic pathway analysis was done using Metacyc database (Caspi et al., 2018).

### QUANTIFICATION AND STATISTICAL ANALYSIS

Statistical analyses of qRT-PCR, chondrocyte metabolites and ketone-glucose blood levels were performed with Excel using unpaired two-tailed Student’s *t*-test. PDH activity assay and metabolic profiling statistics were done using R ANOVA multiple comparison analysis. Statistical significance is denoted by asterisks (P<0.05 [*], P < 0.01 [**], and P < 0.0001 [***]. The data are presented as mean±SD. All statistical details, including n values, are given in the figures and figure legends.

## REFERENCES

1. Harris, R. A., Bowker-Kinley, M. M., Huang, B. & Wu, P. Regulation of the activity of the pyruvate dehydrogenase complex. Adv. Enzyme Regul. 42, 249–259 (2002).

2. Tracy, B. Y., Flora, C. L. & Reed, L. J. a-keto acid dehydrogenase complexes, x. regulation of the activity of the pyruvate dehydrogenase complex from beef kidney mitochondria by phosphorylation and dephosphorylation. 234–241 (1968).

3. O.H., W. The mammalian pyruvate dehydrogenase complex: Structure and regulation. in Reviews of Physiology, Biochemistry and Pharmacology 123–170 (Springer Berlin Heidelberg, 1983).

4. Korotchkina, L. G. & Patel, M. S. Mutagenesis studies of the phosphorylation sites of recombinant human pyruvate dehydrogenase. Site-specific regulation. Journal of Biological Chemistry 270, 14297–14304 (1995).

5. Jeoung, N. H. Pyruvate dehydrogenase kinases: Therapeutic targets for diabetes and cancers. Diabetes Metab. J. 39, 188–197 (2015).

6. P. B. Garland, E. A. Newsholme, and P. J. R. Regulation of Glucose Uptake by Muscle. Biochem. J 93, 665–678 (1964).

7. Kolobova, E., Tuganova, a, Boulatnikov, I. & Popov, K. M. Regulation of pyruvate dehydrogenase activity through phosphorylation at multiple sites. Biochem. J. 358, 69–77 (2001).

8. Bowker-Kinley, M. M., Davis, W. I., Wu, P., Harris, R. A. & Popov, K. M. Evidence for existence of tissue-specific regulation of the mammalian pyruvate dehydrogenase complex. Biochem. J 329, 191–196 (1998).

9. T. E. Roche and Y. Hiromasa. Pdks regulation 2007. Cell. Mol. Life Sci 64, 830–49 (2007).

10. Sugden, M. C. & Holness, M. J. Mechanisms underlying regulation of the expression and activities of the mammalian pyruvate dehydrogenase kinases. (2006). doi:10.1080/13813450600935263

11. Jeong, J. Y., Jeoung, N. H., Park, K.-G. & Lee, I.-K. Transcriptional Regulation of Pyruvate Dehydrogenase Kinase. Diabetes Metab J 36, 328–335 (2012).

12. Semenza, G. L. oxygen regulation semenza 2009. PHYSIOLOGY 24, 97–106 (2009).

13. Aragones, J., Fraisl, P., Baes, M. & Carmeliet, P. Oxygen Sensors at the Crossroad of Metabolism. Cell Metabolism (2009). doi:10.1016/j.cmet.2008.10.001

14. Papandreou, I., Cairns, R. A., Fontana, L., Lim, A. L. & Denko, N. C. HIF-1 mediates adaptation to hypoxia by actively downregulating mitochondrial oxygen consumption. Cell Metab. 3, 187–197 (2006).

15. Kim, J. W., Tchernyshyov, I., Semenza, G. L. & Dang, C. V. HIF-1-mediated expression of pyruvate dehydrogenase kinase: A metabolic switch required for cellular adaptation to hypoxia. Cell Metab. (2006). doi:10.1016/j.cmet.2006.02.002

16. Kluza, J. et al. Inactivation of the HIF-1??/PDK3 signaling axis drives melanoma toward mitochondrial oxidative metabolism and potentiates the therapeutic activity of pro-oxidants. Cancer Res. (2012). doi:10.1158/0008-5472.CAN-12-0979

17. Lu, C. W., Lin, S. C., Chen, K. F., Lai, Y. Y. & Tsai, S. J. Induction of pyruvate dehydrogenase kinase-3 by hypoxia-inducible factor-1 promotes metabolic switch and drug resistance. J. Biol. Chem. 283, 28106–28114 (2008).

18. Prigione, A. et al. HIF1?? modulates cell fate reprogramming through early glycolytic shift and upregulation of PDK1-3 and PKM2. Stem Cells (2014). doi:10.1002/stem.1552

19. Stacpoole, P. W. Therapeutic Targeting of the Pyruvate Dehydrogenase Complex / Pyruvate Dehydrogenase Kinase (PDC / PDK) Axis in Cancer. 109, 1–14 (2017).

20. Tennessen, J. M. et al. Coordinated Metabolic Transitions During Drosophila Embryogenesis and the Onset of Aerobic Glycolysis. doi:10.1534/g3.114.010652

21. Saunier, E., Benelli, C. & Bortoli, S. The pyruvate dehydrogenase complex in cancer: An old metabolic gatekeeper regulated by new pathways and pharmacological agents. International Journal of Cancer (2016). doi:10.1002/ijc.29564

22. Simon, M. C. & Keith, B. The role of oxygen availability in embryonic development and stem cell function. Nat Rev Mol Cell Biol 9, 285–296 (2008).

23. Koh, M. Y. & Powis, G. Passing the baton: The HIF switch. Trends Biochem. Sci. 37, 364–372 (2012).

24. Dunwoodie, S. L. The Role of Hypoxia in Development of the Mammalian Embryo. Developmental Cell (2009). doi:10.1016/j.devcel.2009.11.008

25. Amarilio, R. et al. HIF1alpha regulation of Sox9 is necessary to maintain differentiation of hypoxic prechondrogenic cells during early skeletogenesis. Development 134, 3917–28 (2007).

26. Provot, S. et al. Hif-1?? regulates differentiation of limb bud mesenchyme and joint development. J. Cell Biol. (2007). doi:10.1083/jcb.200612023

27. Schipani, E. et al. Hypoxia in cartilage: HIF-1?? is essential for chondrocyte growth arrest and survival. Genes Dev. 15, 2865–2876 (2001).

28. Hallmann R, Feinberg RN, Latker CH, Sasse J, R. W. Regression of blood vessels precedes cartilage differentiation during chick limb development. Differentiation 34, 98–105 (1987).

29. Maes, C. et al. VEGF-independent cell-autonomous functions of HIF-1?? regulating oxygen consumption in fetal cartilage are critical for chondrocyte survival. J. Bone Miner. Res. 27, 596–609 (2012).

30. Bentovim, L., Amarilio, R. & Zelzer, E. HIF1 is a central regulator of collagen hydroxylation and secretion under hypoxia during bone development. Development 139, 4473–4483 (2012).

31. Vandenboom, R. et al. compensation PDH kinase 2 knockout mice: effect of PDH kinase 1 PDH activation during in vitro muscle contractions in PDH activation during in vitro muscle contractions in PDH kinase 2 knockout mice: effect of PDH kinase 1 compensation. Am J Physiol Regul Integr Comp Physiol Am. J. Physiol. - Regulatory, Integr. Comp. Physiol. Weizmann Inst Sci (2011). doi:10.1152/ajpregu.00498.2010

32. Ho Jeoung, N. et al. Role of pyruvate dehydrogenase kinase isoenzyme 4 (PDHK4) in glucose homoeostasis during starvation. Biochem. J 397, 417–425 (2006).

33. Nam Ho Jeoung, Yasmeen Rahim, Pengfei Wu, W. N. Paul Lee, and R. A. & Harris. Fasting induces ketoacidosis and hypothermia in PDHK2/ PDHK4-double-knockout mice. 443, 829–839 (2012).

34. Semba, H. et al. HIF-1α-PDK1 axis-induced active glycolysis plays an essential role in macrophage migratory capacity. Nat. Commun. 7, 1–10 (2016).

35. Prigione, A. et al. HIF1?? modulates cell fate reprogramming through early glycolytic shift and upregulation of PDK1-3 and PKM2. Stem Cells 32, 364–376 (2014).

36. Provot, S. & Schipani, E. Molecular mechanisms of endochondral bone development. Biochem. Biophys. Res. Commun. 328, 658–665 (2005).

37. Crabb DW, H. R. Mechanism responsible for the hypoglycemic actions of dichloroacetate and 2-chloropropionate. Arch. Biochem. Biophys. 198, 145–152 (1979).

38. Paten, B., Herrero, J., Beal, K., Fitzgerald, S. & Birney, E. Enredo and Pecan: Genomewide mammalian consistency-based multiple alignment with paralogs. 1814–1828 (2008). doi:10.1101/gr.076554.108.

39. Stelzer, G. et al. The GeneCards Suite: From Gene Data Mining to Disease Genome Sequence Analyses. 1–33 (2016). doi:10.1002/cpbi.5

40. Broquist, H. Amino Acid Metabolism. Annu. Rev. Biochem. 35, 231–247 (1966).

41. Stelzer G, Inger A, Olender T, Iny-Stein T, Dalah I, Harel A, Safran M, L. D. GeneDecks: paralog hunting and gene-set distillation with GeneCards annotation. Omi. A J. Integr. Biol. 13, (2009).

42. Szklarczyk, D. et al. The STRING database in 2017: Quality-controlled protein-protein association networks, made broadly accessible. Nucleic Acids Res. 45, D362–D368 (2017).

43. Joshi MA, Jeoung NH, Obayashi M, Hattab EM, Brocken EG, Liechty EA, Kubek MJ, Vattem KM, Wek RC, H. R. Impaired growth and neurological abnormalities in branched-chain α-keto acid dehydrogenase kinase-deficient mice. Biochem. J 162, 153–162 (2006).

44. Logan, M. et al. Expression of Cre Recombinase in the developing mouse limb bud driven by a Prxl enhancer. Genesis (2002). doi:10.1002/gene.10092

45. Stacpoole, P. W. The Pharmacology of Dichloroacetate. Metabolism 38, 1124–1144 (1989).

46. Zhang, S., Zeng, X., Ren, M., Mao, X. & Qiao, S. Novel metabolic and physiological functions of branched chain amino acids: A review. J. Anim. Sci. Biotechnol. 8, 4–15 (2017).

47. Zhou, M., Lu, G., Gao, C., Wang, Y. & Sun, H. Tissue-specific and Nutrient Regulation of the Branched-chain _-Keto Acid Dehydrogenase Phosphatase, Protein Phosphatase 2Cm (PP2Cm) ‪□. 287, 23397–23406 (2012).

48. Nishitani, S. et al. Branched-chain amino acids improve glucose metabolism in rats with liver cirrhosis. 1292–1300 (2019). doi:10.1152/ajpgi.00510.2003.

49. Shao, D. et al. Glucose promotes cell growth by suppressing branched-chain amino acid degradation. Nat. Commun. (2018). doi:10.1038/s41467-018-05362-7

50. Oppenheim, R. D. et al. BCKDH: The Missing Link in Apicomplexan Mitochondrial Metabolism Is Required for Full Virulence of Toxoplasma gondii and Plasmodium berghei. PLoS Pathog. 10, (2014).

51. Wynn, R. M., Chuang, J. L., Cote, C. D. & Chuang, D. T. Tetrameric Assembly and Conservation in the ATP-binding Domain of Rat Branched-chain _-Ketoacid Dehydrogenase Kinase *. 275, 30512–30519 (2000).

52. Gudi, R. et al. Diversity of the Pyruvate Dehydrogenase Kinase Gene Family in Humans. Cell Biol. Metab. 270, 28989–28994 (1995).

53. K M Popov, N Y Kedishvili, Y Zhao, Y Shimomura, D. W. C. and R. A. H. Primary Structure of Pyruvate Dehydrogenase Kinase Establishes a New Family of Eukaryotic Protein Kinases *. J. Biol. Chem. 268, 26602–26606 (1993).

54. Ferriero, R. et al. Phenylbutyrate Therapy for Pyruvate Dehydrogenase Complex Deficiency Phenylbutyrate Therapy for Pyruvate Dehydrogenase Complex Deficiency and Lactic Acidosis. 175, (2013).

55. Huang, B. et al. Isoenzymes of Pyruvate Dehydrogenase Phosphatase. J. Biol. Chem. 273, 17680–17688 (1998).

56. Semenza, G. L. Hypoxia-inducible factor 1: oxygen homeostasis and disease pathophysiology. TRENDS Mol. Med. 7, 345–350 (2001).

57. Kim, J. W., Tchernyshyov, I., Semenza, G. L. & Dang, C. V. HIF-1-mediated expression of pyruvate dehydrogenase kinase: A metabolic switch required for cellular adaptation to hypoxia. Cell Metab. 3, 177–185 (2006).

58. Weissgerber, T. L. & Wolfe, L. A. Physiological adaptation in early human pregnancy: adaptation to balance maternal-fetal demands. 11, 1–11 (2006).

