## Supplemental Information for "BCKDK regulates the TCA cycle through PDC to ensure embryonic development in the absence of PDK family"

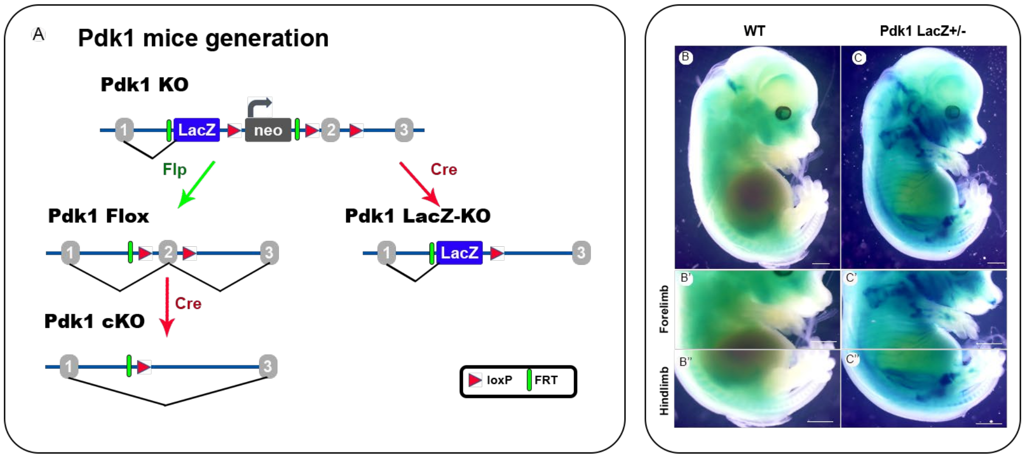


**Fig. S1.** ***Pdk1*** **KO mice generation and its expression in skeletal development**. **Related to Fig. 1** (A) schematic illustration of *Pdk1* strain generation by knockout-first allele method (adapted from EUCOMM) (**B-C''**) X-gal staining of control (B-B") and *Pdk1-lacZ+/-* (C-C") E14.5 embryos demonstrating the expression of *Pdk1* gene in most of the skeletal elements. B',B" and C',C" are magnifications of B and C, respectively. Scale bars: 50 µm in B,C, 100 µm in B'-C''.

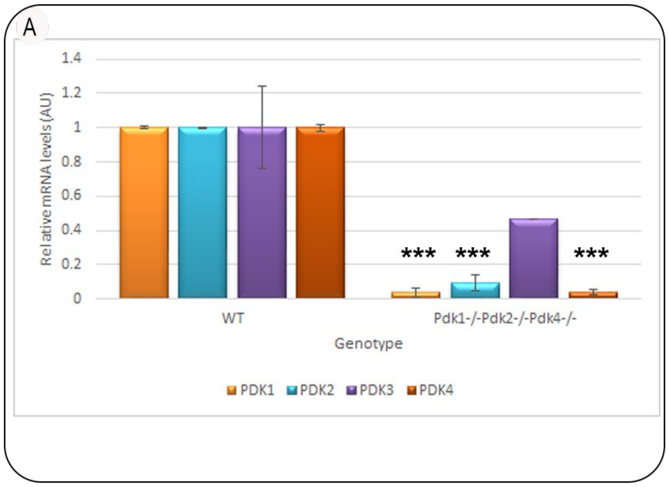


**Fig. S2. *Pdk* tKO validation by mRNA expression**. **Related to Fig. 2 (A)** Graph showing qRT-PCR of *Pdk* mRNA confirms deletion of *Pdk*1, *Pdk*2 and *Pdk*4 genes, as compare to WT (***, $P<0.0001$; $n=3$ for each genotype; data are normalized to *Tbp* and presented as mean±SD).

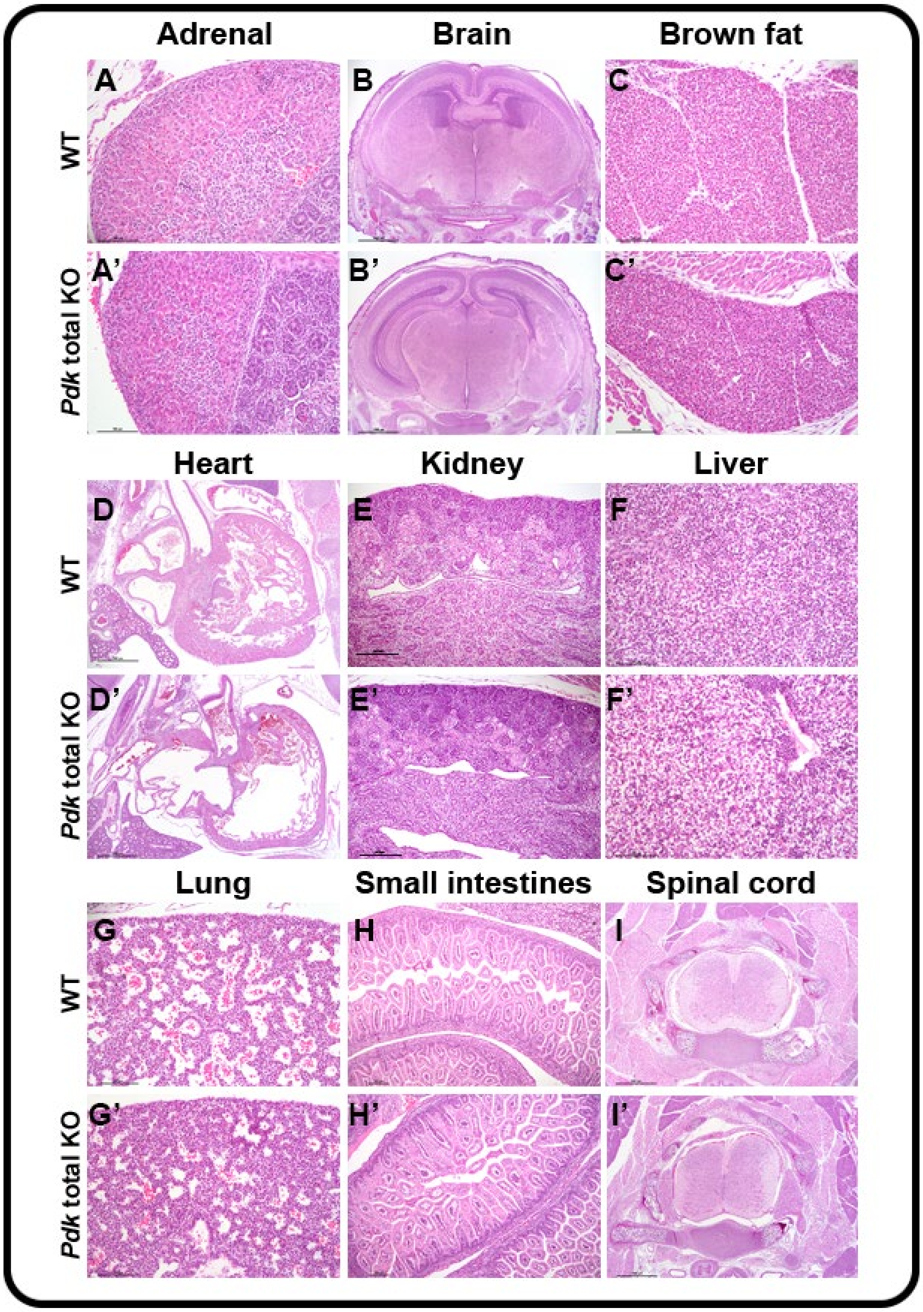


**Fig. S3.** ***Pdk*** **total KO Pathology. Related to Fig. 2**. Pathological examination of E18.5 WT (A-I) and *Pdk* total KO (A'-I') embryos. Representative images of H&E-stained paraffin sections from various organs as indicated ($n_{WT}$=3, $n_{KO}$=3). Scale, 500 µm.

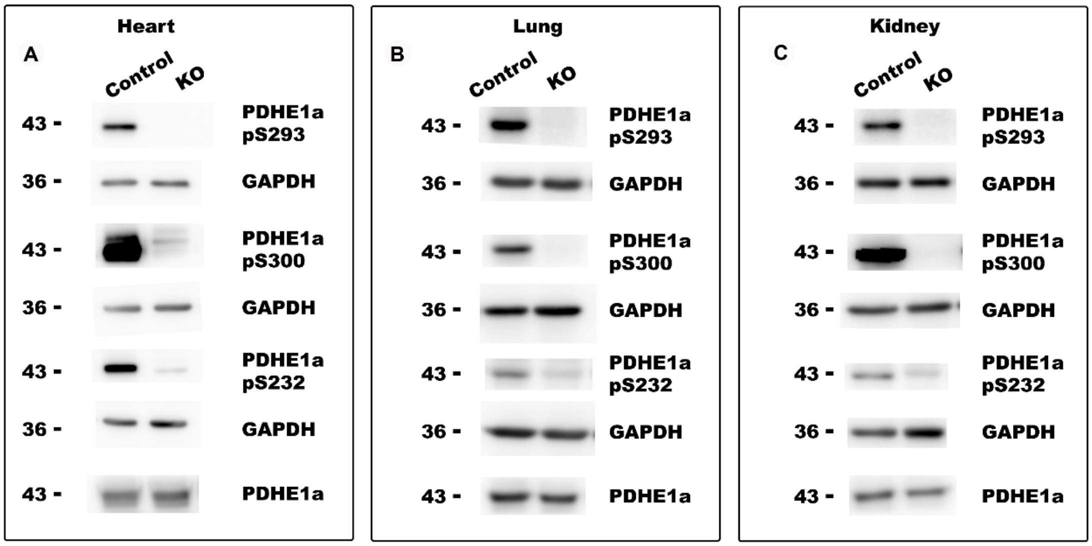


**Fig. S4. PDC phosphorylation in *Pdk* total KO mice. Related to Fig. 3** .Western blot analysis of the three PDC phosphorylation sites in heart, lungs and kidneys extracted from E18.5 *Pdk* total KO embryos, compared to control WT littermates (n=3 for each genotype). As controls, PDHE1a and endogenous GAPDH total protein levels were measured. Total protein concentration loaded was 40 µg.

| **Primer:** | **Sequence:** |
|---|---|
| ***Pdk1* KO / *Pdk1* LacZ forward** | TATTTGTGTCTAGTCAATGACTTGGG |
| ***Pdk1* KO / *Pdk1* LacZ reverse** | CAACGGGTTCTTCTGTTAGTCC |
| ***Pdk1* Flox forward** | TATTTGTGTCTAGTCAATGACTTGGG |
| ***Pdk1* Flox reverse** | AACGGTACATTTACCATAGTGAGAGG |
| ***Pdk2* forward** | TAATCTTGACCCTGGACCAAGG |
| ***Pdk2* reverse WT** | TTGATCTCTTTCATGATGTTGG |
| ***Pdk2* reverse KO** | CGCTTTTCTGGATTCATCGACTGTGGC |
| ***Pdk3* forward** | TTGGAAGCGTAGGACCACAT |
| ***Pdk3* reverse WT** | CCGCGACACCTACACAAGTA |
| ***Pdk3* reverse KO** | TTCAGGAGAGTGCGGTTTGA |
| ***Pdk4* forward** | CTCGAGCGAACACCAATGCACGCTC |
| ***Pdk4* reverse WT** | GGTGCTCGAGCCTGGGTGAAGG |
| ***Pdk4* reverse KO** | CGCTTTTCTGGATTCATCGACTGTGGC |
| ***Bckdk* Forward** | GGAAGACAGGAGCCCTCATAAA |
| ***Bckdk* reverse** | TCTCTGCTGCCACATCAATG |

**Table S1. Genotype PCR primers.**
